# The slow-acting G-protein G_z_ defines the duration of circadian rest time

**DOI:** 10.64898/2026.05.23.727383

**Authors:** Youichi Tanaka, Yuto Fujita, Tom Macpherson, Aoi Harui, Sumihiro Kunisue, Genzui Setsu, Xinyan Shao, Daisuke Ono, Takatoshi Hikida, Yoshihisa Fujita, Michiyuki Matsuda, Hitoshi Okamura, Emi Hasegawa, Masao Doi

## Abstract

Duration of nightly rest is a trait that varies between individuals, influenced by a complex interplay between multiple genetic and environmental factors. The central circadian clock that orchestrates daily rhythm in behavior/sleep resides in the suprachiasmatic nucleus (SCN). Yet, how the SCN encodes the “length” of the circadian rest phase (ρ) remains an open question. Here we demonstrate that the unique G-protein-subtype Gz contributes to this process by sculpting the waveform of the circadian cAMP-PKA activity rhythm within the SCN. Genetic deletion and subsequent rescue of Gz expression reversibly altered the ρ duration, shifting it from ∼10 h in *Gz*^+/+^ mice to ∼7.5 h in *Gz*^-/-;^ mice, accompanied by proportional changes in peak cAMP-PKA activity duration and transcriptome remodeling in the SCN. Notably, intra-SCN cAMP activation led to behavioral rest, with Gz specifically shaping this response without affecting 24-h rhythmicity. These findings suggest that the SCN is not merely a rhythm-generator, but actively allocates the ρ-rest period via Gz signaling.

## INTRODUCTION

The amount of time spent for rest or sleep per night differs between individuals and between animal species— multiple genetic and environmental/social factors influence this trait^1–3^. For example, some people sleep for 5 hours at night in a consolidated block, while others sleep 8 hours; and people keep the same patterns, from day to day, with 24-h rhythmicity. This implies that periodicity and rest duration are likely separate traits. However, we know very little about the mechanism governing the “allocation” of rest time duration in the 24-hour cycle, compared to the well-elucidated circadian clock mechanism responsible for generating 24-h rhythmicity. In mammals, the suprachiasmatic nucleus (SCN) has been defined as the central clock for daily rhythms in behavior and physiology^4,5^. The SCN generates robust 24-h rhythmicity through a mechanism involving cell-autonomous transcriptional feedback control of clock genes as well as inter-cellular coupling via cell-to-cell communication^6^, which ultimately confer 24-h rhythmicity on gene expression as well as cell signaling, metabolism, and neuronal firing rate in the SCN^6–8^. Nevertheless, the mechanism by which the SCN allocates circadian rest timing and duration in each 24-hour cycle remains largely unknown. The manners in which rest timing and duration are expressed at the molecular level in the SCN are not fully understood.

The unique G-protein subclass Gz (also initially referred to as Gx) is an evolutionarily conserved vertebrate G-protein that belongs to the family of canonical Gi/o (ref^9,10^). Similar to Gi1, Gi2, and Gi3, Gz inhibits the activity of adenylyl cyclases and thereby reduces cAMP synthesis^11^. However, Gz is not just an “ersatz” Gi. As its name suggests, Gz displays a range of unique properties that differentiate it from other Gi/o members. First, in contrast to the ubiquitous expression of Gi1/2/3, tissue distribution of Gz is rather specific to brain^12,13^ including the SCN^14^. Secondly, the Gi/o inhibitor, pertussis toxin (PTX), does not inhibit Gz. PTX catalyzes ADP ribosylation at a specific cysteine residue conserved in Gi/o. The absence of this residue in Gz makes it resistant to PTX, posing hurdles in pharmacological approach to this protein^15^. Thirdly, and equally as important as the first two, Gz bears relatively low GTPase activity^16^. The *k*_cat_ values for GTP hydrolysis by Gi/o and Gs are in the range of 1–5 min^−1^; however, Gz displays an activity approximately 100 times slower (*k*_cat_, ∼0.05 min^−1^)^15,16^. Thus, once GTP binds to it, Gz does not readily return to its inactive form, resulting in a prolonged signal in terms of temporal resolution, which suggest differential roles for Gz and other Gi in brain signal processing. Lastly, only scanty information is currently available for the function of Gz in CNS: previous studies demonstrated that Gz knockout mice exhibit altered neuronal and neuroendocrine responses to several pharmacological treatments such as those with morphine^17^, dopamine D2 receptor agonist^18^, and serotonin receptor 5-HT1A agonist^19,20^; however, its endogenous function in brain’s physiology remains to be explored, including its contribution to the circadian clock.

In the present study, we report the identification of a critical contribution of Gz to the mechanism of “rest time allocation”. The genetic deletion and subsequent restoration of Gz expression in the SCN reversibly altered the circadian rest bout length, shifting it from ∼10 h in *Gz*^+/+^ mice to 7.5 h in *Gz*^-/-;^ mice (here, the “*rest*” time represents the interval from the activity offset to the next onset in each circadian cycle^21,22^). *Gz* deficiency did not affect 24-h rhythmicity of the circadian clock, reinforcing the idea that Gz specifically contributes to the trait of rest time allocation. At the molecular level, we explored the contribution of Gz to cAMP signaling and protein kinase A (PKA) activity in the SCN. To address this molecular signaling and further define the causal relationship between Gz and rest–time allocation, we performed fiber-photometry-based simultaneous monitoring of behavior and PKA activity rhythm in the SCN, using freely moving mice (see Methods). Our data show that Gz-mediated signaling is required for the maintenance of the shape of cAMP-PKA activity rhythm in the SCN.

## RESULTS

### Antiphasic 24-hour coordination of SCN PKA activity and behavior in freely moving mice

The SCN cAMP signal fluctuates in a circadian fashion in in-vitro cultured SCN slices^23,24^. In the present study, we monitored the downstream PKA activity oscillation, in vivo. To determine the precise temporal relationship between the day/night rest–activity profiles and in vivo SCN circadian PKA activity rhythms over multiple days, we used fiber photometry (Fig. 1a, schematic cartoon). In this experiment, we used a genetically encoded FRET-based PKA biosensor, AKAR3EV (ref^25^) for monitoring PKA activity in vivo and we employed infrared video tracking system (ref^26^) to simultaneously monitor animal daily behavior (Fig. 1a).We conducted long-term recording, initially under a regular 12-h light:12-h dark cycle (LD) condition and then constant dark (DD) condition, followed by a light pulse stimulation in animal’s subjective night (Fig. 1b). Notably, real-time monitoring revealed consistently anti-phasic relationships between in vivo FRET activity in the SCN and organismal spontaneous behavioral activity, under LD-entrainment condition, DD free-running condition, and even after a light pulse-induced phase shift of the rhythms (see double-plotted temporal profiles of circadian FRET activity in *green* and behavioral activity in *blue*, as well as horizontal serial temporal profiles in Fig. 1b). This antiphasic temporal organization of PKA activation and behavior is consistent with previous observations from time-series cAMP enzyme-immunoassay^27^ as well as cAMP signal-related immunohistochemistry^28,29^, but is more precise than these assays in terms of time resolution (every 5 min) and monitoring length (e.g. 16 days in Fig. 1b; see also Supplementary Fig. 1).

**Fig. 1.**
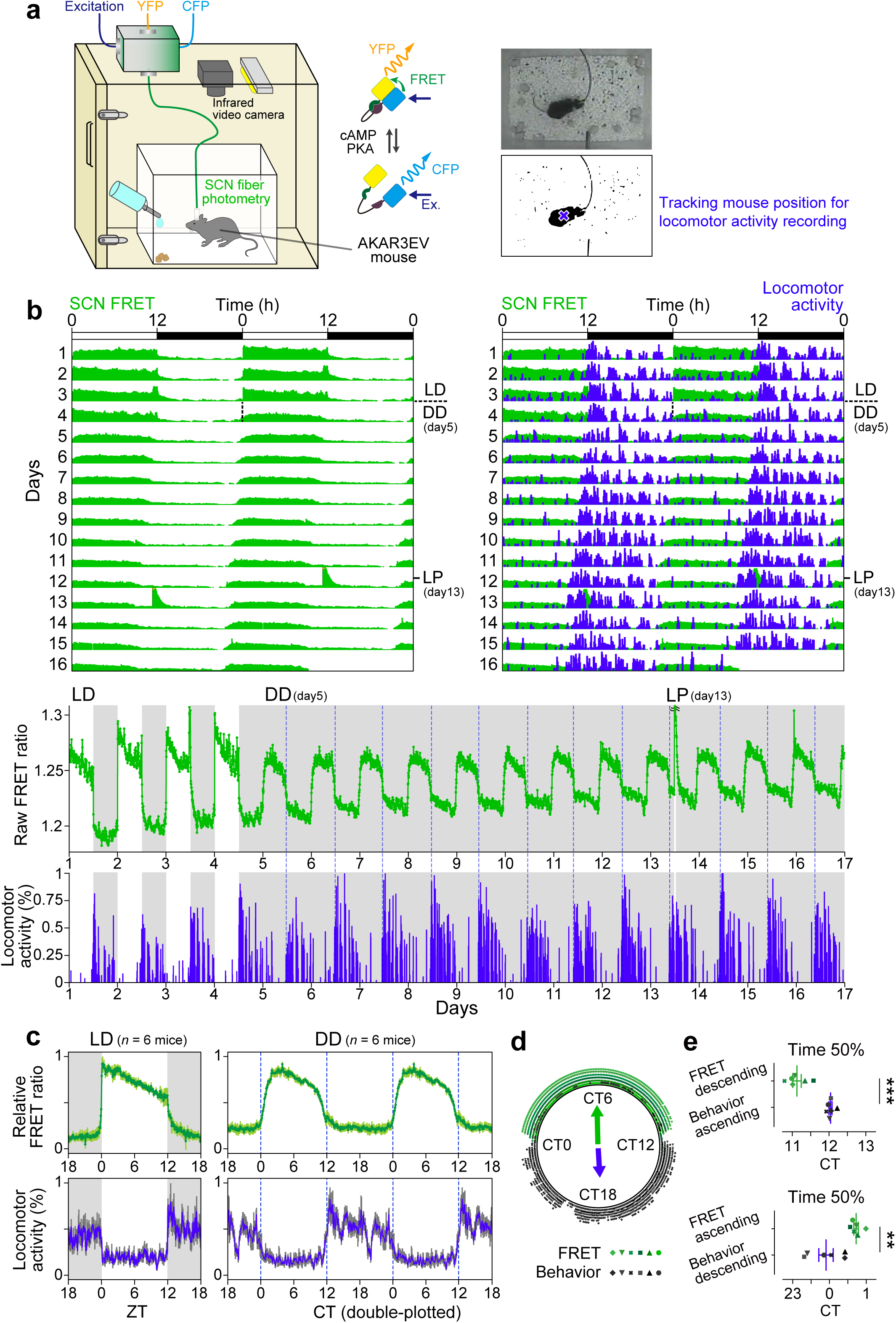
Longitudinal monitoring of in vivo SCN PKA activity and behavioral rhythms in freely moving mice. **a** Experimental setup. AKAR3EV, a FRET-based biosensor for PKA activity, was used to trace SCN PKA activity via fiber photometry. Simultaneously, locomotor activity was monitored via an infrared camera. Pictures show example video images in the light and dark period. **b** Representative double-plotted SCN PKA FRET activities (left). Mice were placed in LD from day 1 to day 4 and thereafter kept in DD from day 5 to day 16, except for receiving a brief light pulse (LP) at CT14 on day 13. Spontaneous locomotor activity (blue) was overlaid on the FRET activity (green) record to allow direct phase comparison (right). The same data were also plotted horizontally in line (bottom) (periods of darkness are indicated by shaded grey backgrounds). **c** Averaged daily activity profiles of SCN FRET and behavior under LD and DD conditions. *n* = 6, AKAR3EV mice. Error bars indicate s.e.m. **d** Circular Rayleigh plot showing phase distribution of SCN FRET activity (green) and behavior (black). The data points above the mean of total FRET/behavior activity are plotted with symbols representing different animals (*n* = 6). Arrows in the circle are the Rayleigh plot vector. **e** Quantification of the 50% ascending and descending times for FRET activity and behavior (**d**) around CT12 and CT0. \*\*\**P* < 0.001, \*\**P* < 0.01. Paired two-sided Student’s *t*-test.

The shape of rhythm, revealed by our assay, attracted our attention in characterizing the SCN PKA FRET activity rhythms. Firstly, we noted that the shape of the PKA FRET activity rhythms in the SCN was not in a smooth sinusoidal waveform. The inactive (thus, nadir) period of the FRET PKA activity rhythms in the SCN was almost flat, forming a broad and extended trough. Moreover, we found a noticeable light-dependent difference in the shape of rhythm (waveform) between LD and DD (Fig. 1c, LD vs. DD): the FRET rhythms in LD exhibit a steep rising at the beginning of the light period (by definition, at zeitgeber time (ZT) 0) and a sharp decreasing at the end of the light period (ZT12); while in DD the corresponding changes were modest. The rhythm waveform, thus, adopted a “square” pattern under LD cycles, while displaying a bell-shaped curve in DD (Fig. 1b,c).

When mice underwent eye enucleation, the SCN FRET activity free-ran with a rhythm-shape of bell type (DD type) without acute FRET changes at the transitions of L/D, excluding the possibility of ambient light-induced artificial noise affecting our recording (Supplementary Fig. 1a). Of note, a light pulse given at CT14 in the subjective night (to non-enucleated mice) led to a high-amplitude induction of the FRET ratio in the SCN, and this was followed by a phase shift of the FRET rhythms in the cycles after the light pulse (LP, Fig. 1b)(see also Supplementary Fig. 1b). Importantly, enucleation caused abolishment of both light-induced acute upregulation of FRET ratio and phase shift (sham operation vs. enucleation: FRET fold change relative to the circadian peak level, 3.57 ± 0.63 vs. –0.01 ± 0.02; phase delay, 1.29 ± 0.15 h vs. 0.01 ± 0.02 h; mean ± s.e.m., *P* < 0.01, two-sided *t*-test, Supplementary Fig. 1c,d). Thus, the light-dependent shape of SCN PKA activity was a result of retinal input, instead of an artifact photometry.

The chief advantage of dual monitoring of intra-SCN activity and behavioral outcome is its ability to trace the exact mutual temporal relationship (Fig. 1c-e), a question of neuroethology. The antiphasic oscillations of SCN PKA activity and behavior were constantly observed in DD with an almost identical free-running period, both slightly less than 24 h (23.74 ± 0.04 h for FRET, 23.80 ± 0.10 h for behavior, determined by FFT-NLLS; *n* = 6). We plotted averaged daily profiles of these two rhythms in the same graph to allow comparison of their temporal relationship (Fig. 1c; CT12 was defined based on behavior for each animal; see Methods for detail). We found that the start of rising of FRET activity in the SCN corresponds to the start of the behaviorally inactive (sedentary) period. The end of waning of the SCN FRET activity, on the other hand, corresponds to the onset of nocturnal behavioral activity (CT12, by definition) (Fig. 1d). We found that during the sedentary period, when animals were quiescent and their behavior at basal levels, FRET activities were maintained at mid-to-high levels. Notably, the FRET activities began decreasing, a few hours before behavioral onset. The time at 50% reduction of FRET activity (CT11.13) was earlier than that of half-maximal induction of behavioral activity around CT12 (CT12.14) (see Fig. 1e, *P* < 0.001, paired *t*-test). These data suggest that the mid-to-high, sustained acrophase of intra-SCN PKA activity is related to the maintenance of rest duration.

### The G-protein Gz deficiency causes shortening of rest period and PKA acrophase length

With the aim of identifying a relevant molecular mediator of the circadian PKA activity in the SCN, we focused on studying the G-protein Gz (Fig. 2). Gz is a member of the heterotrimeric Gi protein family with predominant expression in the brain, including the SCN^12–14^, but its role in the SCN, particularly its contribution to circadian behavioral regulation in vivo, is not known. We found that mice with a targeted deletion of the gene encoding the α-subunit of Gz (*Gnaz*), referred to here as *Gz*^-/-;^ mice (Fig. 2)^17^, display a significantly shortened circadian rest period compared to wildtype control mice (*Gz*^+/+^), and concomitantly display SCN PKA activity with limited peak activity duration (Fig. 2a–e, behavior; Fig. 2f–i, FRET PKA activity; see below for quantification of data).

**Fig. 2.**
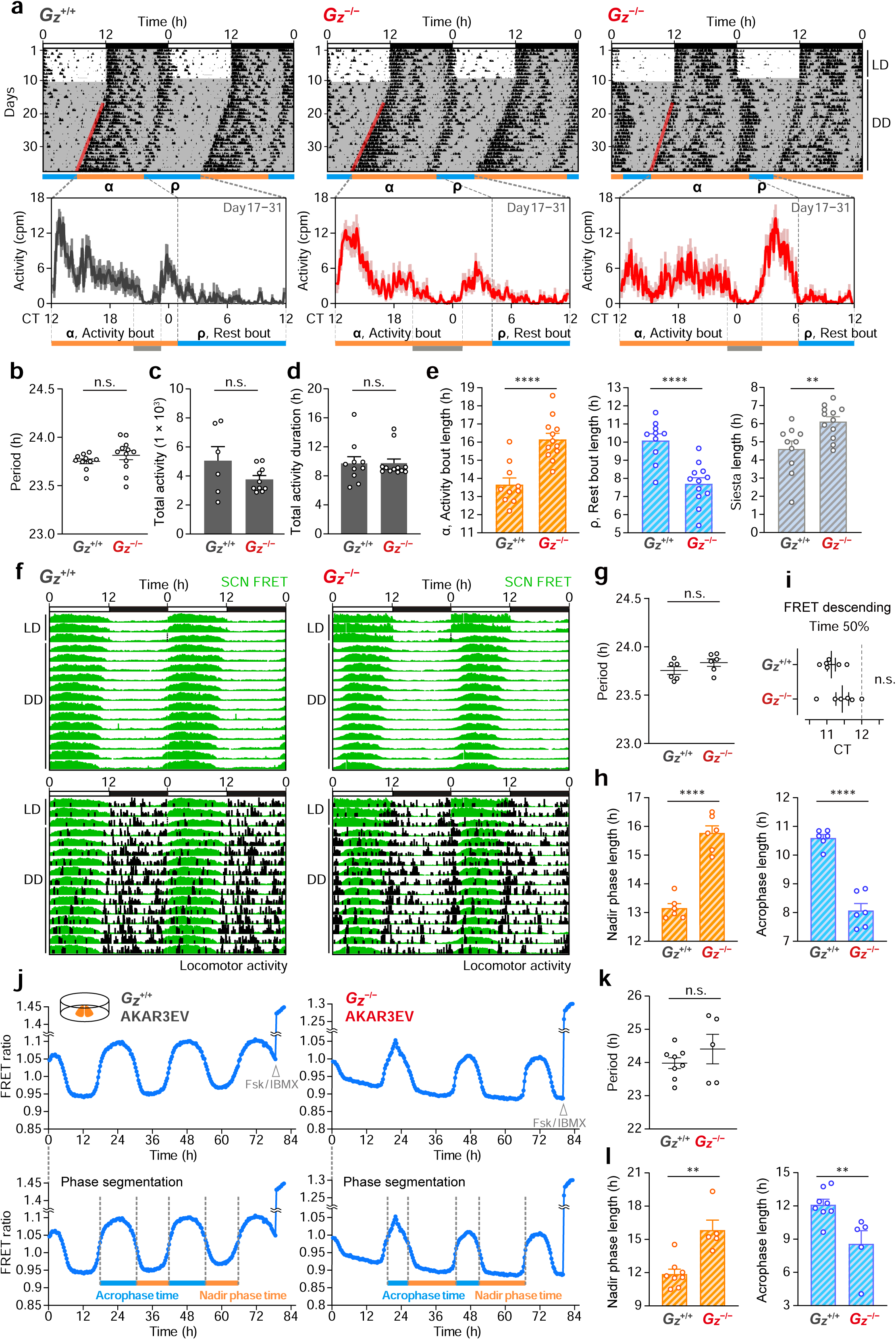
Effects of *Gnaz* deficiency on circadian behavior and SCN-PKA activity profiles. **a** Representative double-plotted locomotor activity plots of *Gz*^+/+^ and *Gz*^-/-;^ littermate mice housed in LD followed by DD. Red lines delineate the phase of activity onset. Averaged circadian activity profiles starting from CT12 are displayed in line graphs. The colored horizontal bars indicate differential activity bout (α) and rest bout (ρ) lengths, determined by the offset of before-dawn locomotor activity. The grey bars depict a siesta in the middle of the activity period. **b-e** Quantification of the period length (**b**), total activity (**c**), total activity duration (**d**) and α, ρ and siesta length (**e**) of *Gz*^+/+^ mice (*n* = 10) and *Gz*^-/-;^ mice (*n* = 12). \*\**P* < 0.01, \*\*\*\**P* < 0.0001. **f** Representative double-plotted *in vivo* SCN-AKAR3EV FRET activity plots of *Gz*^+/+^ and *Gz*^-/-;^ mice, overlaid with their spontaneous locomotor activity, in LD followed by DD. **g-i** Quantification of period length (**g**), nadir and acrophase length (**h**), and the 50% descending time (**i**) of *in vivo* FRET activity rhythms in (**f**). *n* = 6, AKAR3EV *Gz*^+/+^ mice; *n* = 6, AKAR3EV *Gz*^-*/*-^ mice. \*\*\**P* < 0.001. **j** Representative *ex vivo* AKAR3EV FRET activity traces from cultured SCN slices of *Gz*^+/+^ and *Gz*^-/-;^ mice. The traces below contain plots showing nadir phase (orange) and acrophase (blue) separation. Fsk/IBMX was applied for slice viability. **k**, **l** Quantification of period length (**k**) and nadir phase and acrophase length (**l**) of (**j**). Period was determined by fast Fourier transform–nonlinear least squares analysis (FFT-NLLS). *n* = 8, AKAR3EV *Gz*^+/+^; *n* = 5, AKAR3EV *Gz*^-*/*-^ SCN slices. \*\**P* < 0.01. Error bars in (**b-e**, **g-i**, **k**, **l**) indicate s.e.m. Statistics, unpaired two-sided Student’s *t*-test. n.s., not significant.

First, based on the actograms, we compared the parameters of behavior (*ρ*, *α*, and *τ*) of *Gz*^-/-;^mice and *Gz*^+/+^ mice: Nocturnal activities in both mice were bimodal, showing morning and evening components consistent with previous reports^21,22,30,31^. However, a genotype-related behavioral difference was evident in the duration of “rest” time (*ρ*): here, as per the definition, *ρ* represents the interval from the activity offset to the next onset in circadian cycles (see *blue* bars beneath the actograms, which represent the *ρ* bout length, Fig. 2a)^22^. Quantification revealed the *ρ* of *Gz*^-/-;^ mice being approximately 2.5 hours shorter than that of *Gz*^+/+^ mice (*P* < 0.0001, two-sided *t*-test, Fig. 2e; *ρ*, 7.66 ± 0.36 h for *Gz*^-/-;^ mice, *n* = 12; 10.09 ± 0.36 h for *Gz*^+/+^, *n* = 10). Notably and consistently, SCN PKA activity measured by FRET in *Gz*^-/-;^ mice displayed a similar contraction of the duration of acrophase PKA activity (see Fig. 2f): when the FRET acrophase was defined as the interval between the mid-timepoint of rising phase and the mid-timepoint of decreasing phase in each cycle, we found the shorter acrophase of *Gz*^-/-;^ mice by approximately 2.5 h relative to *Gz*^+/+^ mice (8.07 ± 0.25 h for *Gz*^-/-;^; 10.59 ± 0.12 h for *Gz*^+/+^; *P* < 0.0001, two-sided *t*-test, *n* = 6, both genotypes, Fig. 2h). A similar phenotype was also expressed by SCN slice cultures, in vitro (Fig. 2j–l): *Gz*^-/-;^ SCN slices recapitulated circadian FRET PKA activity rhythms with limited peak duration—and its peak duration was noted as approximately 3.5 h shorter than that of *Gz*^+/+^ slices (*Gz*^+/+^, 12.11 ± 0.47 h; *Gz*^-/-;^, 8.57 ± 1.09 h; Fig. 2l); Thus, the phenotype is inherent to the SCN; and rather the exacerbated phenotype was observed in vitro. Our data indicate that *Gz* deficiency led to the shortening of PKA acrophase in the SCN and ρ in behavior, in mice.

At the expense of decreased *ρ* length, “activity” time (*α*, the length between activity onset and offset as per the definition^21^) was accordingly increased in *Gz*^-/-;^ mice by approximately 2.5 h (*Gz*^+/+^, 13.66 ± 0.36 h; *Gz*^-/-;^, 16.15 ± 0.33 h, Fig. 2e; see also the orange *α* length in Fig. 2a). Nevertheless, this did not indicate hyperactive phenotype: Although the time length defined as *α* was lengthened, there was a compensatory decrease in activity level in the middle of *α* period defined in *Gz*^-/-;^ mice (see the rest zone, *grey* in Fig. 2a), which results in no statistical difference between *Gz*^-/-;^ mice and *Gz*^+/+^ mice in either the total activity amount (Fig. 2c) or total activity duration (Fig. 2d) per day. Moreover, we observed that the values of free-running period (*τ*)—the sum of *α* and *ρ*—were almost identical between the two genotypes, meaning that *Gz*^-/-;^ mice are normal in terms of periodicity (c.f. Fig. 2b for behavior; Fig. 2g,k, PKA in vivo and in vitro activity). Our data therefore indicate that whereas *Gz*^-/-;^ mice displayed a severe contraction of the SCN PKA activity acrophase and rest bout length, these two events maintain their mutual synchrony with a period of approximately 24 h (see coordinate anti-phasic FRET activity, *green*, and behavior, *black*, in *Gz*^-/-;^, and 50% FRET descending, prior to behavioral onset, for *Gz*^-/-;^, Fig. 2f,i). This close co-occurrence of abnormal peak PKA activity length and diminished *ρ* (Fig. 2f) suggests that the main direct behavioral phenotype of *Gz*^-/-;^is causing diminution of *ρ*, without affecting τ.

We observed that actograms of all examined *Gz*^-/-;^ mice display *ρ* shortening (Supplementary Fig. 2). Using electroencephalogram (EEG), we extended this observation. Overall, *Gz*^-/-;^ mice were normal in spectral distribution of EEG power, total time, episode number and episode duration for wake, NREM, and REM sleep in the entire 24-hour day (Fig. 3a,b, telemetry data). However, the length of consolidated circadian rest period, determined by sleep–wake cycle (i.e., duration between circadian sleep onset and offset) was shortened in *Gz*^-/-;^ mice (7.73 ± 0.33 h) compared to *Gz*^+/+^ mice (10.40 ± 0.28 h) (*P* < 0.001, *ρ*, *Gz*^-/-;^ vs. *Gz*^+/+^, two-sided *t*-test; *n* = 6 mice for *Gz*^-/-;^; *n* = 4 mice for *Gz*^+/+^) and a compensatory increase in siesta sleep was observed for *Gz*^-/-;^ mice (Fig. 3c,d) without affecting *τ* (*Gz*^+/+^, 23.62 ± 0.06 h, *Gz*^-/-;^, 23.64 ± 0.11 h) and the total amount of sleep time (see Fig. 3d). These data are compatible with the actogram-based data (Fig. 2a–e) and reinforce the phenotype of reduced *ρ* duration in *Gz*^-/-;^ mice.

**Fig. 3.**
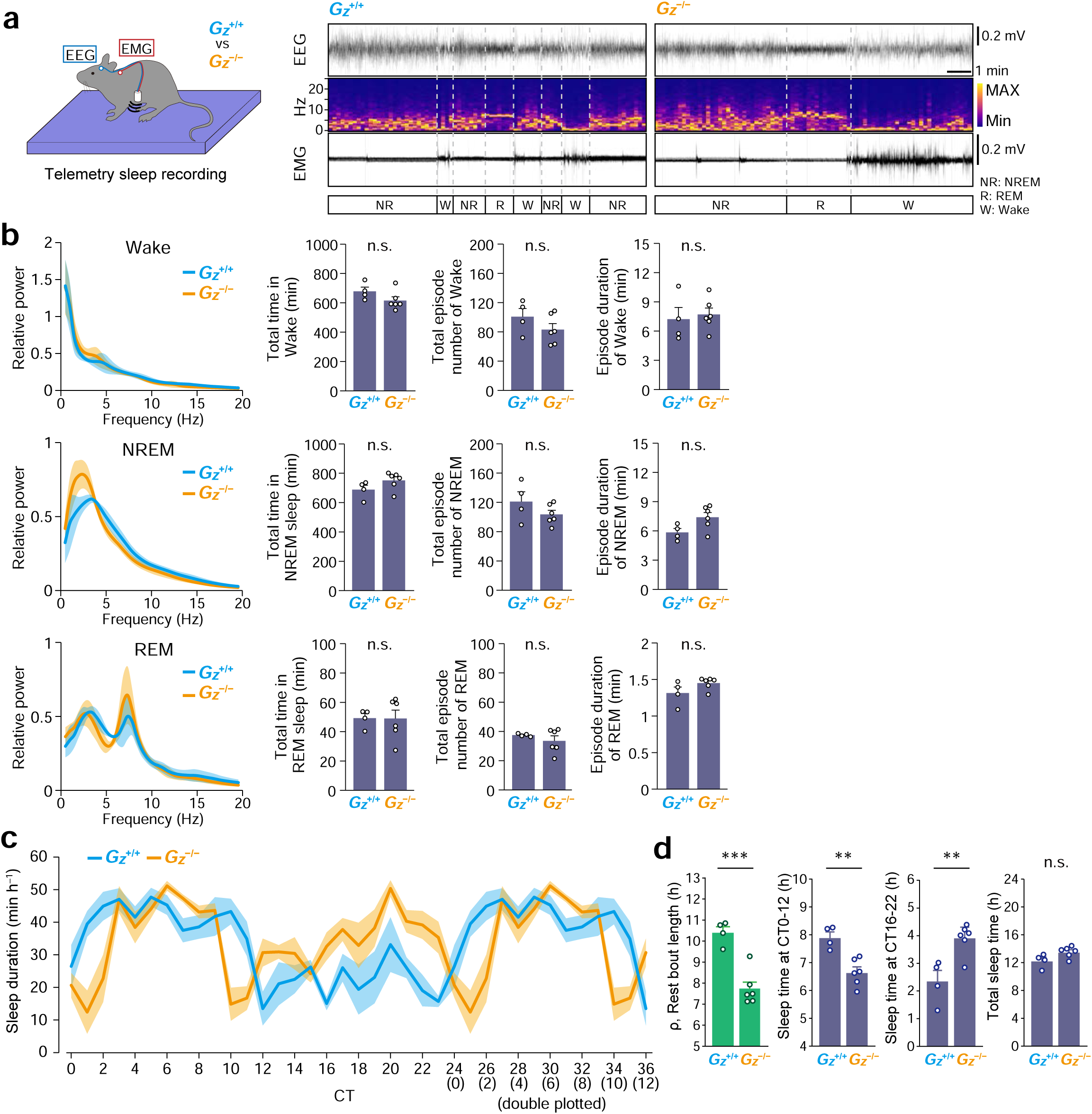
Assessment of circadian rest phase duration via sleep−wake cycle recording. **a** Schematic of telemetry sleep recording and representative EEG traces, EEG power spectra, EMG traces, and assigned vigilance states of *Gz*^+/+^ and *Gz*^-*/*-^ mice. **b** EEG power spectra during wakefulness, NREM, and REM sleep in *Gz*^+/+^ (*n* = 4) and *Gz*^-*/*-^ mice (*n* = 6). Values (mean ± sd) show data from 3 consecutive circadian cycles. Bar graphs indicate the averaged total time, episode number, and duration of wakefulness, NREM, and REM sleep per circadian cycle. **c** Circadian profiles of hourly sleep duration of *Gz*^+/+^ and *Gz*^-*/*-^ mice (**b**). Data are double plotted. Light shaded areas depict s.e.m. **d** Quantification of ρ length and sleep time at CT0-12, CT16-22, and CT0-24 (total). CT6 was presented as the midpoint of ρ. \*\*\**P* < 0.001, \*\**P* < 0.01, two-sided *t*-test. Error bars within the bar graphs of (**b**) and (**d**) indicate s.e.m.

### Re-expression of *Gz* in the SCN is sufficient to reverse *ρ* length in *Gz*^-/-;^ mice

Since *Gz* mRNA expression is not restricted to the SCN and rather broadly distributed across the brain^32^, *Gz* deletion in extra-SCN regions might account for the phenotype of *Gz*^-/-;^ mice. It is also possible that the *ρ* phenotype could arise secondarily from a potential developmental defect of SCN due to the absence of *Gz* during development. We thus next sought to examine whether re-expressing *Gz* in the SCN in adult mice has the effect of reversing the *ρ* phenotype of *Gz*^-/-;^ mice (see Fig. 4). To test this, we monitored *ρ* of mice before and after viral injection of adeno-associated virus (AAV) vector encoding either *Gz* (AAV-hSyn-Gz-2A-mCherry) or mCherry only (as a control: AAV-hSyn-mCherry) to the SCN of *Gz*^-/-;^ mice (Fig. 4a). Before treatment, individual *Gz*^-/-;^ mice were confirmed to show a shortened circadian *ρ* length of about 8 hours (7.92 ± 0.41 h, Fig. 4b). Notably, however, the *ρ* of *Gz*^-/-;^ mice that underwent AAV-mediated *Gz* reexpression in the SCN was lengthened to 11.82 ± 0.44 h (*P* < 0.0001, paired *t*-test vs. pre-AAV, *n* = 11, Fig. 4b) and this value somewhat surpasses that observed in wildtype *Gz*^+/+^ mice (10.09 ± 0.36 h, see Fig. 2e). In contrast to this marked *ρ* extending effect, re-expression of *Gz* did not significantly alter the circadian period (*τ*), with individual variations smaller than ∼0.3 h (see Fig. 4b). Importantly, all examined individual *Gz*^-/-;^ mice that received *Gz* reexpression in the SCN displayed a reversal of *ρ* without notable change in *τ* (individual actogram data are available in Supplementary Fig. 3). On the other hand, *Gz*^-/-;^ mice that received AAV injection of control mCherry vector (AAV-hSyn-mCherry) in the SCN continued showing the *ρ* shortening phenotype (*P* = 0.203, pre-vs. post-AAV, *n* = 9, Fig. 4c; see also Supplementary Fig. 3) with almost unaffected *τ* length as expected. These data indicate that delivered *Gz* expression but not *mCherry* in the SCN is effective and sufficient to reverse the *ρ* phenotype of *Gz*^-/-;^ mice. This effective rescue of *ρ* by *Gz* within the SCN echoes the idea that the SCN is a central organizer for circadian behavioral control.

**Fig. 4.**
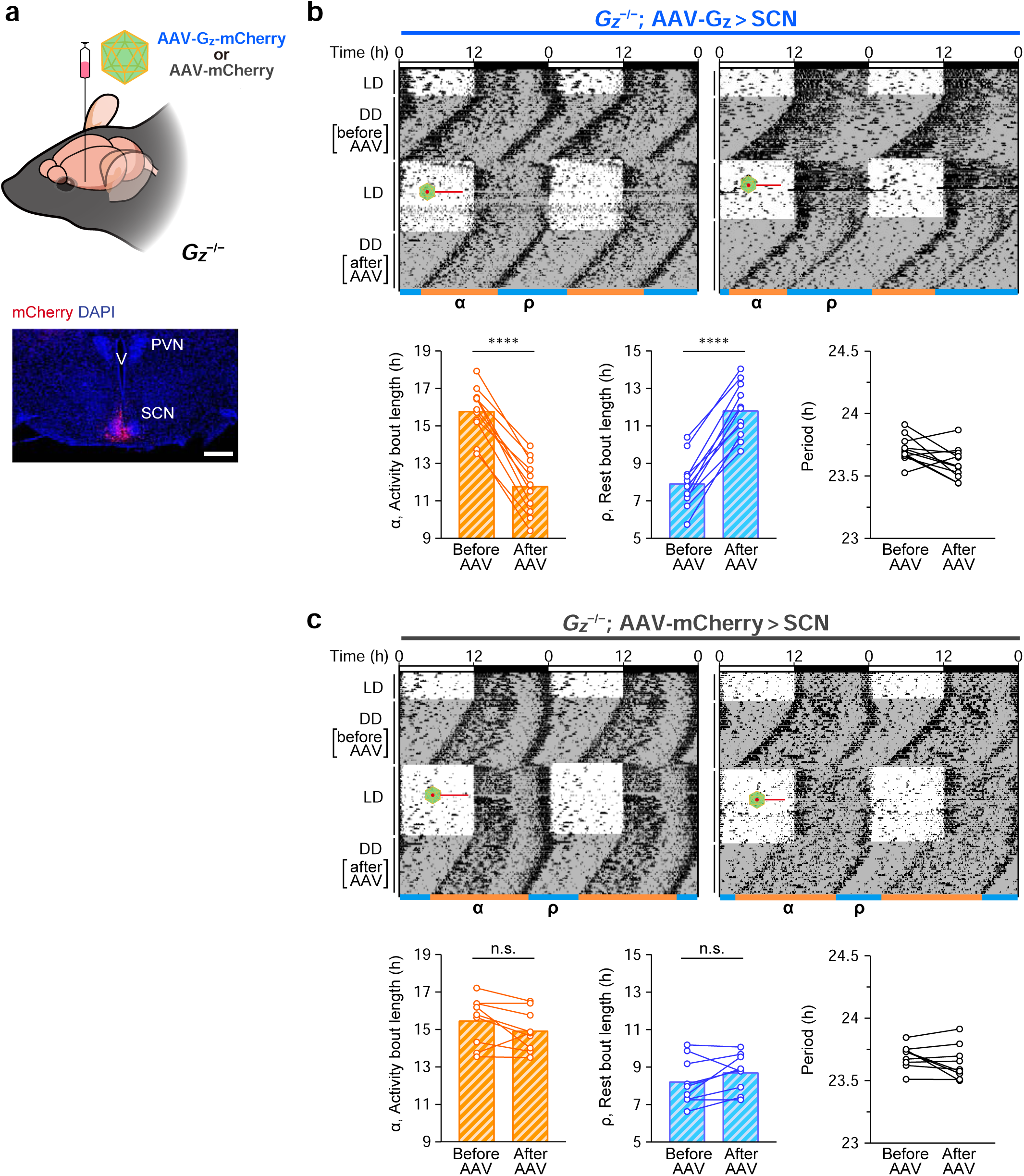
Re-expression of Gz in the SCN reconstitutes normal rest bout length in *Gz*^-/-;^ mice. **a** Injection of AAV-Gz-2A-mCherry or its control AAV-mCherry into the SCN of *Gz*^-/-;^ mice. The brain section shows mCherry fluorescence in the SCN of AAV-injected mice. Scale bar, 0.4 mm. **b** Representative double-plotted actograms and quantification of α, ρ, and period length of *Gz*^-/-;^ mice before and after AAV-Gz-2A-mCherry injection (*n* = 11 mice). Red lines indicate the time of operation for viral injection. **c** Same as (**b**) but for AAV-mCherry injection (*n* = 9 mice). \*\*\*\**P* < 0.0001. Paired two-sided *t*-test. n.s., not significant.

### *Gz* expression remodels the shape of autonomous circadian PKA-activity rhythms in the SCN

Our data suggest that re-expression of *Gz* can remodel or restore the SCN function – even after development. To pursue this implication, we next used SCN slice cultures, with two questions: 1) Can the *Gz* reexpression restore the aberrant shape of PKA activity in in vitro SCN slices? —this question regards whether the steady state (or established rhythm shape) of PKA activity in the slices (autonomous in vitro rhythms) can be remodeled after *Gz* reexpression—; and 2) What temporal transitions underlie this remodeling process?—given that Gz inhibits cAMP signaling, how the shape (or equilibrium) shifts upon *Gz* reexpression remains a question.

We traced the shape of circadian FRET PKA activity rhythms before and after infection of *Gz* or *mCherry* to cultured organotypic SCN slices (Fig. 5). Before treatment, the rhythms of *Gz*^-/-;^slices (see Fig. 2j–l) were characterized by abnormally elongated nadir phase duration. This elongation of the nadir period was, however, somehow counterintuitive at the molecular level because the ablation of Gz, a cAMP inhibitory factor, would reduce the inhibitory (nadir) period rather than extend it. Resolving this, we observed that re-expression of *Gz* reduced the nadir basal levels of PKA activity, resulting in a sharper and narrower trough, within 3–5 days post AAV infection (Fig. 5a,c,d). The control expression of *mCherry* did not produce such changes, importantly: Therefore, it is *Gz* but not *mCherry* or infection artifact that caused the shape of PKA-activity rhythm to change. Quantification of our data verified that the *τ* was unaffected by *Gz* reexpression (Fig. 5b), while the values of the nadir-bottom length as well as the nadir mean intensity of PKA activities were reduced significantly post *Gz* expression (Fig. 5c, nadir length, pre-AAV, 11.93 ± 0.81 h, post, 9.00 ± 1.15 h; Fig. 5d, nadir intensity, relative to pre-AAV, 0.972 ± 0.004 at the 3rd–5th cycles, *n* = 7 for both conditions).

**Fig. 5.**
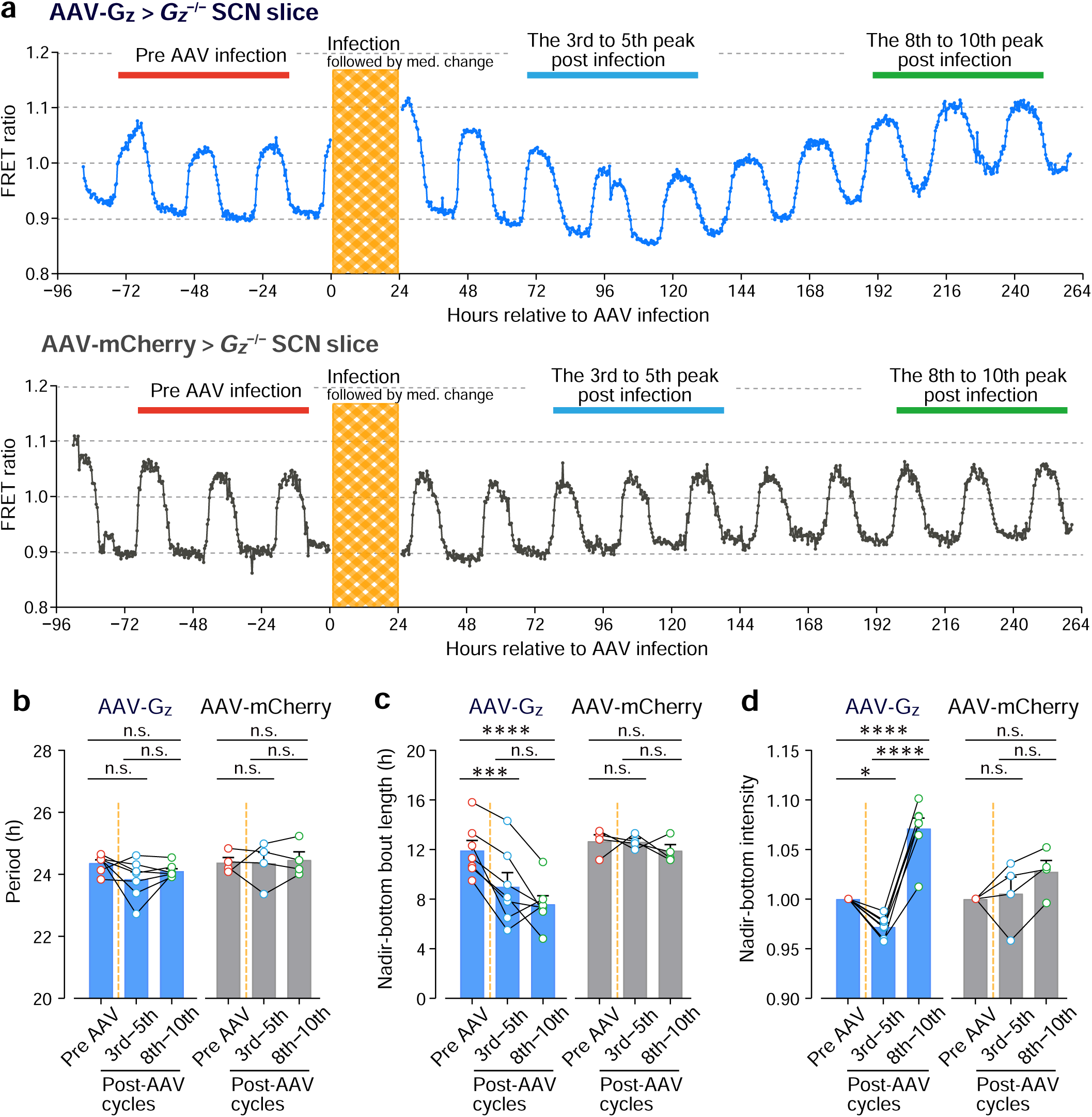
Re-expression of Gz reforms trough PKA activity. **a** Representative AKAR3EV FRET activity traces from cultured *Gz*^-/-;^ SCN slices infected with AAV-Gz-2A-mCherry (upper) and AAV-mCherry (lower). The shaded zone indicates the period of AAV treatment. **b-d** Quantification of nadir-bottom bout length (**c**), nadir-bottom intensity (**d**), and period length (**b**) of the FRET activity rhythms before and after (selected cycles) AAV infection. The data represent biologically independent slices, *n* = 7 for AAV-Gz-2A-mCherry and *n* = 4 for AAV-mCherry. \*\*\*\**P* < 0.0001, \*\*\**P* < 0.001, \**P* < 0.05. Error bars indicate s.e.m. Repeated measures two-way ANOVA with Bonferroni’s *post hoc* test. n.s., not significant.

Based on the phenotypes obtained at 3–5 days post infection, we expected a continuation of similar rhythms with reduced nadir activity, aligning with the expected role of *Gz*. However, unexpectedly, when the cultures were kept for an extended period (∼8–10 days), we observed that the levels of PKA activity ceased decreasing and turned to go up to the levels even higher than those of pre-AAV infection, for all rescued slices tested (Fig. 5a,d, nadir intensity, relative to pre-infection, 1.07 ± 0.01 at the 8th–10th cycles, *P* < 0.0001, repeated measures two-way ANOVA with Bonferroni’s test). This rising was associated with a continuation of a shortened nadir bout length (*P* = 0.22, the 3rd–5th vs. the 8th–10th cycles, Fig. 5c). These data suggest that the remodeling process of PKA activity rhythms due to *Gz* reexpression in the SCN may involve multiple unknown regulatory processes that independently influence the shape and level, which cannot be explained by a simple cAMP-inhibitory action of *Gz*.

### *Gz* expression upregulates the cAMP signaling activator *Gs* expression

To gain insight into the mechanisms underlying the up-shifting of PKA activity rhythms in *Gz* rescued SCN slices, we conducted RNA-seq analysis (Fig. 6). After 10 days post-infection, the *Gz*^-*/*-^ SCN slices infected with *Gz* or control *mCherry* were collected (*n* = 3 RNA-seq samples for each group; each was generated from a pool of 3 slices). For consistency, all rescued slices were confirmed by FRET monitoring to have elevated PKA activity rhythms before sampling. We collected the samples at the early acrophase on day 10. Furthermore, to ensure the elimination of potential confounding factors derived from infection, we concomitantly analyzed the RNA samples prepared from non-infected wildtype *Gz*^+*/+*^ slices as well as non-infected *Gz*^-*/*-^ slices for comparison (Fig. 6a, *left*, non-infected *Gz*^+*/+*^ vs. *Gz*^-*/*-^ SCN; *right*, AAV-*Gz* rescue vs. AAV-control *Gz*^-*/*-^ slice transcriptome). This combinatorial, nonbiased analysis revealed that the gene encoding the stimulatory G-protein Gs α-subunit (*Gnas*) —a stimulator of cAMP synthesis—was included among the genes that are upregulated upon rescue of *Gz* expression (Fig. 6b, *Gnas*). *Gnas* expression was oppositely decreased by the absence of *Gz*, suggesting a balance between Gs and Gz (inhibitory) for homeostatic cAMP coordination. This associated expression of *Gnas* could partly explain the mechanism of up-shifting of PKA activity due to *Gz* expression.

**Fig. 6.**
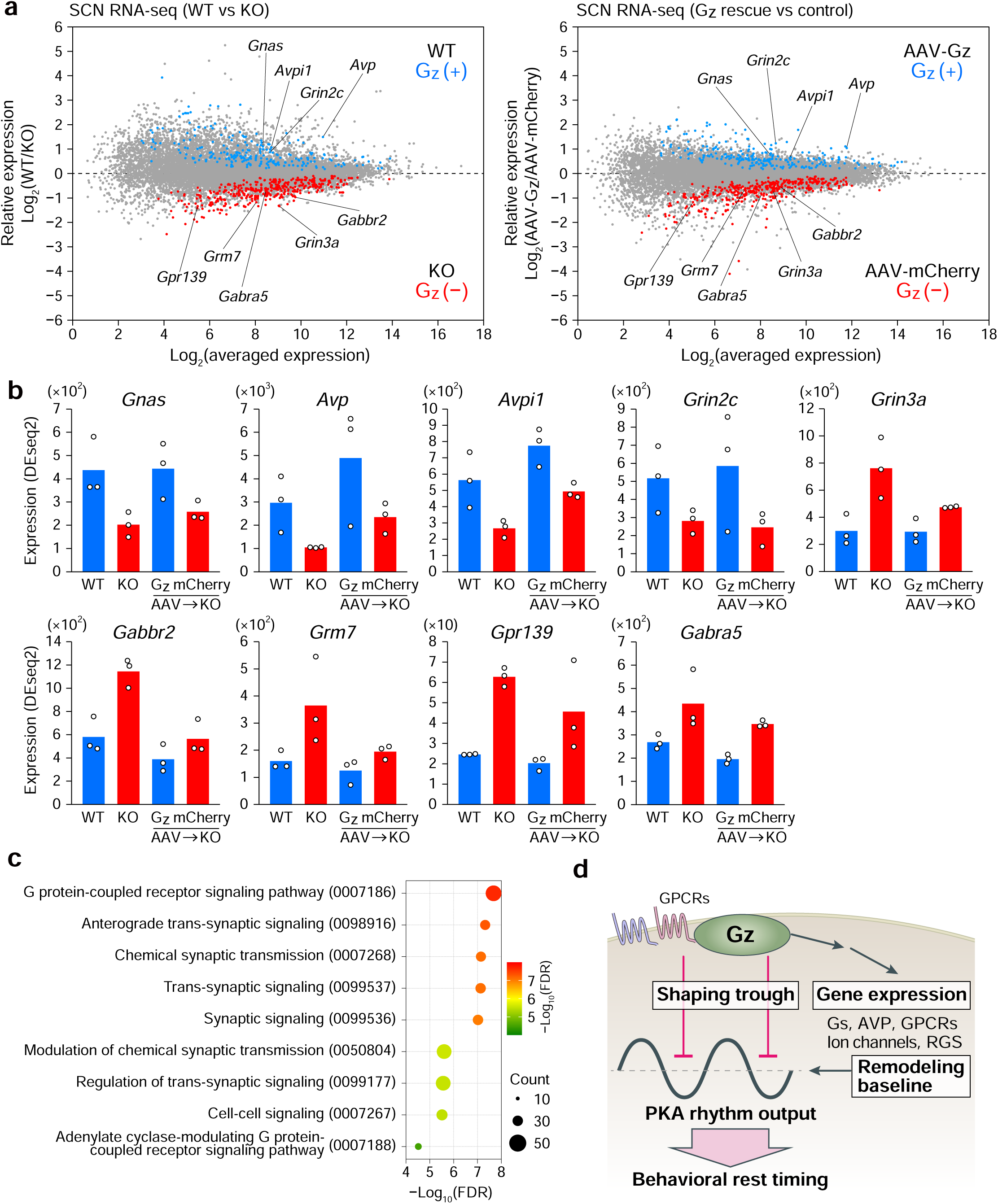
Gz expression remodels gene expression of cAMP signaling mediator in the SCN. **a** RNA-seq data showing WT versus *Gz*^-*/*-^ SCN slice transcriptome (*left*) and AAV-Gz rescue versus AAV-control *Gz*^-*/*-^ slice transcriptome (*right*), presented in MA-plot format. Values are based on the mean of *n* = 3 biologically independent RNA-seq samples for each group; each RNA-seq sample was generated using a pool of 3 SCN slices. Plots in blue and red represent transcripts significantly upregulated and downregulated in Gz (+) conditions, respectively (*P*_adj_ < 0.05, DESeq2 two-way ANOVA Gz (+) vs Gz (−)). **b** Representative gene transcripts annotated as upregulated or downregulated in Gz (+) conditions in (**a**). Note that the transcript encoding cAMP activator Gnas (Gsα) was decreased in *Gz*^-*/*-^ (KO, red) compared to *Gz*^+*/+*^ (WT, blue) and restored by AAV-based Gz rescue (blue) compared to mCherry control (red). **c** Bubble plot for GO enrichment analysis based on Gz (+) vs Gz (−) conditions. **d** Schematic model depicting a direct and indirect role of Gz in shaping circadian PKA activity in the SCN.

Besides *Gs*, there was a widespread transcriptomic remodeling after *Gz* re-expression in *Gz*^-*/*-^slices. Therefore, we compared *Gz*^-*/*-^ vs. *Gz*^+*/+*^ and AAV-*Gz* rescue vs. AAV-control *Gz*^-*/*-^ SCN transcriptome and generated groups of genes affected by the presence of *Gz* (*blue* highlights genes commonly up-regulated by the presence of *Gz*, *red* commonly up-regulated by absent *Gz*) (Fig. 6a). Gene Ontology (GO) enrichment analysis of differentially expressed genes (Fig. 6a, *blue*, 203 and *red*, 259; the gene lists are available in Supplementary Data 1) revealed the highest enrichment in the “pathway of G protein coupled receptor signaling”, involving over 46 genes, including *Gabbr2* (GABA B receptor 2, known to signal to Gi), *Grm7* (Glutamate metabotropic receptor 7, coupled to Gi), *Gpr139* (GPR139, orphan), and *Gnas* (Fig. 6b, 6c, Supplementary Data 1). The ion channel-coding genes, *Gabra5* (GABA A receptor subunit alpha 5), *Grin2c* (Glutamate [NMDA] receptor subunit 2C) and *Grin3a* (Glutamate [NMDA] receptor subunit 3A) were also differentially expressed according to *Gz*-expression (Fig. 6b). We also observed that arginine vasopressin (*Avp*) gene, known to function as a neuropeptide crucial to maintain proper circadian oscillation of the SCN^33^, and its downstream gene, *Avpi1* (Arginine vasopressin-induced 1), were down-regulated in *Gz*^-*/*-^ slices and restored upon *Gz* reexpression (Fig. 6b). These adaptive and reversible gene expression changes indicate that *Gz* gene is a functional component affecting the state of gene-expressions in the SCN. Besides acting as a direct modulator of cAMP signaling leading to functional PKA activity fluctuations, Gz therefore likely exerts its indirect effects leading to expressional changes of G-protein Gs, neuropeptide AVP, other GPCRs, ion channels, and RGS. Our data suggest that, whether direct or indirect, or a combination of both, the actions of rescued *Gz* in the SCN ultimately reshape circadian PKA activity levels and resultant acrophase length in the SCN (Fig. 6d; model).

### Validation of circadian signaling tracked by PKA FRET activity

Finally, we wanted to validate our method. We have used our PKA-sensor as a Gz-cAMP signal reader reflecting the central clock’s circadian activity. However, this may not be true. A potential caveat to the results of our FRET PKA activity monitoring (in Fig. 1−6) is that it may unintentionally reflect cAMP-unrelated or circadian clock-independent signal variations in the SCN, which could lead to misleading observations. To address this concern, we finally examined the FRET activity patterns in SCN slices lacking the core clock component *Bmal1* (*Bmal1*^−/−^, Fig. 7a,b). We also examined wildtype slices under pharmacological inhibition of adenylyl cyclase (AC) activity (Fig. 7c,d). *Bmal1*^−/−^ slices produced low-amplitude, unstable rhythmicity, with periods varying 13 to 20 h—significantly shorter than the consistent ∼24 h of wildtype slices. Unstable peak-to-peak intervals were also notable in *Bmal1*^−/−^ slices (Fig. 7a,b). Pharmacological inactivation of AC with a potent, selective inactivator MDL-12,330A suppressed circadian FRET PKA activity rhythms in wildtype SCN slices immediately after administration, while FRET activity recovered promptly upon removal of the inhibitor (Fig. 7c,d). These data confirmed that the oscillations of PKA activity observed thus far are driven by the circadian clock and are lying downstream of AC-mediated signaling. To directly assess the cAMP rhythm, we also adopted a luciferase-based cAMP biosensor, Okiluc-aCT (ref^24^). AAV encoding Okiluc-aCT was transduced into SCN slice cultures (Fig. 7e, cartoon). This assay, importantly, reproduced the elongated nadir/shortened acrophase phenotype of *Gz*^-*/*-^SCN slices (nadir length, *Gz*^+/+^, 12.42 ± 0.48 h, *Gz*^-/-;^, 15.23 ± 0.53 h, *P* < 0.01; acrophase, *Gz*^+/+^, 12.40 ± 0.63 h, *Gz*^-/-;^, 9.28 ± 0.70 h, *P* < 0.01; Fig. 7e,f)—about 3 h difference between genotypes, without affecting total *τ,* validating our FRET PKA activity observations (Fig. 2l). Lastly, we manipulated intracellular cAMP using a *Beggiatoa* photoactivatable AC (bPAC)^34^ and asked whether this cAMP signal activation in the SCN has a role in reducing behavioral activity. As shown in Fig. 7g, we virally introduced bPAC expression in the SCN using AAV hSyn-flex-bPAC vector in *Slc32a1-Cre* mice for SCN targeting^24,35^. We noted that optogenetic stimulation of cAMP synthesis led to reduced behavior, while the same optic stimulation in control mice expressing only mCherry had no significant effect on behavior (*P* < 0.01, only for pre-vs. during stimulation in bPAC-expressing mice). These data are consistent with our model (Fig. 6d) and explain the antiphasic coordination between cAMP/PKA activity rhythm and behavior in vivo (Fig. 1,2).

**Fig. 7.**
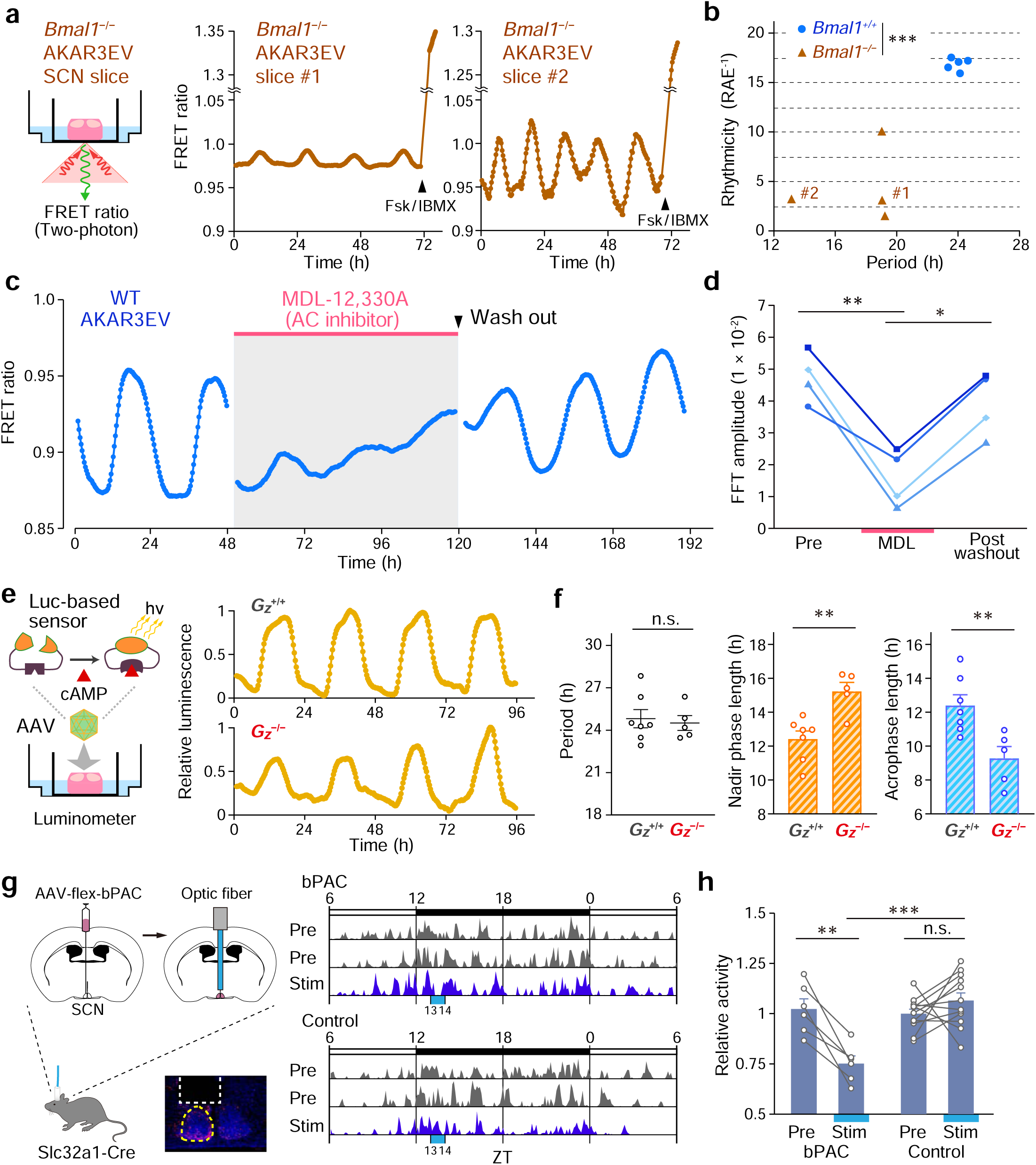
Circadian PKA activity arising from SCN intrinsic clock-mediated cAMP signaling. **a** Representative records of AKAR3EV FRET activity fluctuations in *Bmal1*^-*/*-^ SCN slices. Fsk/IBMX was applied for validation of two-photon imaging and SCN slice viability. **b** FFT-NLLS analysis of rhythms in (**a**) and WT slices, shown as period-versus-rhythmicity plot for WT (*n* = 5) and *Bmal1*^-*/*-^ (*n* = 4 biologically independent slices). Rhythmicity is calculated as the reciprocal of relative amplitude error (RAE^-1^). \*\*\**P* < 0.001. **c** Reversible suppression of circadian AKAR3EV FRET activity by MDL-12,330A treatment. **d** FFT-NLLS amplitude analysis of (**c**). Each data line represents a single SCN slice (*n* = 4). \**P* < 0.05, \*\**P* < 0.01. **e** Representative bioluminescence traces from cultured *Gz*^+/+^ and *Gz*^-/-;^ SCN slices infected with AAV-Okiluc-aCT. **f** Quantification of period, nadir phase, and acrophase length of (**e**). *n* = 7, *Gz*^+/+^; *n* = 5, *Gz*^-*/*-^ Okiluc-aCT SCN slices. \*\**P* < 0.01. **g** Actograms of mice before and after upregulation of SCN cAMP signaling by introduced bPAC optic activation. **h** Quantification of behavioral activity in (**g**). The activities during optic activation of SCN cAMP at ZT13–14 (Stim) were compared with those during the corresponding ZT in preceding two days (Pre). Error bars indicate s.e.m. Statistics, multivariate ANOVA with Wilks’ lambda test for (**b**), one-way ANOVA followed by Bonferroni’s test for (**d**), unpaired two-sided *t*-test for (**f**), repeated measures two-way ANOVA with Bonferroni’s *post hoc* test for (**h**). n.s., not significant.

## DISCUSSION

Daily profiles of rest–activity behavior are expressed as an output of the master clock residing in the SCN. Besides providing periodicity (*τ*) to the organism, the SCN has a role in directing timing and duration of behavior – “active (*α*)” versus “rest (*ρ*)” – of the organism. However, contrasting to the extensive research on the rhythm-generation mechanism, the manners in which daily temporal patterns of *ρ* (or *α*) are expressed at the molecular level in the SCN in vivo have remained unexplored. By combination of animal behavior monitoring and long-term fiber photometry-based real-time tracing of PKA activity in freely moving mice, we revealed that the circadian PKA activity that the SCN displayed in vivo occurred synchronously with *ρ*. Of note, the unique G-protein subclass Gz contributes to this process as a key regulator for cAMP-PKA signaling in the SCN. The genetic deletion and subsequent restoration of Gz expression reversibly alter the circadian rest (*ρ*) bout length, shifting it from ∼10 h in *Gz*^+/+^ mice to 7.5 h in *Gz*^-/-;^ mice, accompanied by proportional changes in the PKA acrophase length in the SCN. This reversibility only affects the length of *ρ* and peak duration of SCN cAMP-PKA activity rhythm, without affecting circadian period (*τ*). Thus, Gz is uniquely suited in the central clock “output” system governing *ρ*: Our data show that Gz contributes to the definition of circadian rest period of the organism by sculpting the waveform of circadian cAMP-PKA activity rhythm in the SCN (see our model summary in Supplementary Fig. 4).

Our study was conducted at the intersection of Gz research field and circadian clock biology. Gz has been a mysterious G-protein subtype, uniquely characterized by its brain-specific expression, PTX insensitivity, low intrinsic GTPase activity compared to other Gi/o, lack of information of downstream transcriptional effect, and exclusive conservation within vertebrates (e.g., absent in flies). These characteristics have historically hindered in-depth functional analyses of this G-protein subtype. In this context, we have provided evidence that Gz acts as a functional G-protein contributing to hourly regulation of cAMP pathway in the SCN. Compared to canonical Gi/o, Gz has significantly lower GTPase activity—approximately 100 times slower (*k*_cat_ values for GTP hydrolysis by Gi/o are in the range of 1–5 min^−1^, whereas Gz ∼0.05 min^−1^)^15,16^. As a result, once activated (GTP-bound), Gz remains in its active state for an extended duration, leading to a prolonged signal. This characteristic may be relevant for Gz to participate in hourly regulation of circadian clock. Our RNA-seq data further identified Gz’s downstream transcriptomic effect—an indirect role of Gz signaling—which may contribute to the homeostatic control of the basal level of cAMP signaling. Particularly, Gz reversibly altered expression of *Gs*, indicating the importance of balance of Gs (“accelerator”) and Gz (“brake”) for equilibrium. Although flies do not possess Gz (ref^9^), the ancestral G-protein Gi likely takes on a counter-balancing role with Gs in the fly’s pacemaker neurons, where the Gs-coupled pigment dispersing factor (Pdf) receptor PdfR and the Gi-coupled metabotropic glutamate (Glu) receptor mGluRA are reported to control circadian cAMP signal oscillation in a cooperative manner^36,37^. Relatedly, our RNA-seq data identified Grm7, a homolog of mGluRA, as being upregulated by Gz deficiency (Fig. 6), implying a common mechanism shared by species whereas *Gz* itself is known as a vertebrate innovation^9,10^.

We showed that re-expression of *Gz* can restore the SCN function — even after development and even after isolation in culture (Fig. 4, behavior; Fig. 5, SCN in culture). These observations strongly suggest that all molecular underpinnings required for the *Gz*-mediated restoration are present in the SCN even after development and even after isolation—without any additional or supplemental input from regions outside the SCN. We therefore surmise that Gz-linked GPCRs that can function in the local SCN network may be involved in this tissue-autonomous remodeling mechanism. One may speculate a potential involvement of Gpr176, a Gz-linked GPCR present in the SCN^14^; however, *Gpr176*^-/-;^ mice do not show *ρ-*shortening phenotype^38^. A comprehensive understanding of Gz-linked GPCR signal in the SCN is therefore required for the elucidation of the mechanism of *ρ* restoration by *Gz*—a direction of our future study. Among other GPCRs in the SCN, serotonin 5-HT_1A_ receptor couples with both Gz and Gi. A potential involvement of *Gabbr2* and *Grm7* (two GPCRs identified as DEG in *Gz*^-/-;^ SCN; Fig. 6b) is also worth investigation as they might be able to couple with Gz in addition to Gi.

We took advantage of dual real-time monitoring of intra-SCN activity and behavior. This allowed us to recognize the unique temporal profile of in vivo PKA activity in the SCN (Fig. 1). A conventional view is that the time-course of cAMP-PKA activation is rather transient; yet it does persist for over ten hours in the SCN in normal wildtype mice. Moreover, the timing of this activity is strictly aligned with the occurrence of *ρ*. Regardless of the genotype (*Gz*^+/+^ or *Gz*^-/-;^), the rise and fall of SCN PKA activity coincide with the initiation and termination of *ρ*, so that the PKA activity acrophase is tightly allocated to *ρ* in each cycle. These hourly-scaled unique temporal profiles likely represent the feature of the SCN output flow, regulating ρ. The inactive period of the PKA activity in the SCN was almost flat, forming a broad and extended trough, which further supports its role in defining the boundaries between α and ρ. These observations and implications may form a basis of neuroethology for understanding the circadian dynamics of rest–activity behavior.

How do our findings relate to research on sleep regulation?–It has been proposed that two key processes underlie the regulation of sleep, one represented by homeostatic sleep regulation and the other by circadian control on sleep timing (see refs^39–41^). In this paradigm, the role that we identified for Gz in the SCN—namely, the determination of the time and duration of consolidated circadian rest by Gz—can be ascribed to the circadian process. However, we should mention that *Gz* is not necessary for either rhythm-generation or period-determination; our data suggest that Gz plays a role in a unique niche within the circadian process as it only affects the size of ρ (output shaping: Supplementary Fig. 4). It is worth noting that the total sleep time per day of *Gz*^-/-;^ mice is equivalent to that of wildtype mice (Fig. 3b), indicating that homeostatic sleep regulation is not impaired in these mice. *Gz*^-/-;^ mice exhibit increased sleep time in the middle of *α* (Fig. 3c), suggesting a compensatory siesta rest in these mice. These data reinforce the idea that Gz contributes to the circadian process without compromising sleep homeostasis. A potential functional crosstalk between the processes of circadian ρ length regulation and homeostatic sleep regulation is an intriguing possibility that remains to be studied.

How the “length” of consolidated block of sleep/rest time zone is determined is an issue independent of how it cycles in a certain periodicity. In this research background, by studying the previously uncharacterized G-protein subtype Gz (Gx), we have obtained the first molecular entity that contributes to the allocation of ρ. Gz reversibly modulates the length of ρ without affecting *τ*. Thus, the SCN is not simply a 24-h metronome. In the SCN, Gz constitutes the mechanism of time allocation of specific output behavior —ρ— by sculpting the waveform of cAMP-PKA activity rhythm in the tissue. We therefore demonstrate that the SCN encodes the information of *ρ* length via a signal through Gz. Given the reversible effect of *Gz* on the duration of *ρ*, modulating Gz-mediated signaling—either activation or inhibition — may have potential implications for *ρ*-rest time intervention.

## STAR METHODS

### RESOURCE AVAILABILITY

#### Lead contact

Further information and requests for resources and reagents should be directed to and will be fulfilled by the lead contact, Masao Doi.

### Material availability

This study did not generate new unique reagents. All materials, including AAV vector constructs, are available through requests to the corresponding authors.

### Data and code availability

- RNA-seq data have been deposited at GEO (accession # GSE284123) and are publicly available as of the date of publication. Accession numbers are listed in the key resources table. All other data reported in this paper will be shared by the lead contact upon request.
- This paper does not report original code.
- Any additional information required to reanalyze the data reported in this paper is available from the lead contact upon request.

## EXPERIMENTAL MODEL AND SUBJECT DETAILS

### Animals

C57BL/6J *Gzα-*deficient (*Gnaz*^−/−^) mice were generated by Hendry *et al*.^17^ and obtained from the Australian Phenome Bank (C57BL/6JAnu-Gnaz^tm1Iah^/AnuApb). AKAR3EV mice (https://animal.nibn.go.jp/e_pkachu.html) were bred as described^42–45^ and intercrossed with *Gnaz*^−/−^ mice^17^ or *Bmal1*^−/−^ mice^46,47^. *Slc32a1-Cre* mice were obtained from the Jackson Laboratory (*Slc32a1^tm2(cre)Lowl^*/J: #016962). Mice were reared at 22–23 °C with a 12 h light/12 h dark cycle and *ad libitum* access to food and water. For circadian locomotor activity analysis, adult mice (8-12-week old) were housed individually in light-tight, ventilated cabinets. Unless otherwise mentioned, behavioral activities were monitored via passive infrared sensors (FA-05F5B; Omron) and analyzed with ClockLab software (Actimetrics), as previously described^48^. All animal experiments were conducted in compliance with ethical regulations in Kyoto University and performed under protocols approved by the Animal Care and Experimentation Committee of Kyoto University.

### Organotypic SCN slice culture

Organotypic SCN slices were prepared as previously described^49^ with slight modifications. In brief, brains from neonatal (3-7-day-old) AKAR3EV mice were coronally sliced into 400-µm-thick slices with a McIlwain tissue chopper, and slices containing the SCN were further dissected under a stereomicroscope to minimize the presence of regions outside the SCN. The resultant SCN explants (approximately 0.7 mm long and 0.7 mm wide) were cultured on a Millicell membrane insert (PICM0125, Millipore) using a medium containing 50% minimum essential medium, 25% Hank’s balanced salt solution, 25% horse serum, 36 mM D-glucose and, antibiotics at 35 °C for at least 2 weeks before FRET imaging. We used a Millicell insert whose plastic struts beneath the membrane had been removed. This ensures the specimen to be placed within the depth-of-field of the FRET microscope we used (LCV110-MPE, Olympus).

## METHOD DETAILS

### In vivo fiber photometry recording

At the beginning of surgery, mice were anesthetized with 3% isoflurane and positioned in a stereotaxic apparatus (Kopf). Anesthesia was maintained with 1.5–2.0% isoflurane for the duration of the surgery. Lidocaine (8%) was administered topically on the scalp. A midline scalp incision was made, and the skull was cleaned in order to visualize bregma and lambda. The angle of the head was adjusted so that the skull was flat. A small burr hole was made at AP, −0.4 mm, ML, 0.0 mm from the bregma using a high-speed micro drill (Foredom). Then, an optical fiber was implanted (0.5 NA, 400 μm diameter, Kyocera) 0.2 mm above the SCN. Dental resin (Metabond, Parkell) was used to securely attach the fiber optic cannula to the skull. The area of attachment was coated with black opaque dental cement to avoid light contamination. Animals were allowed to recover for 2 weeks before experiments proceeded.

For fiber photometry, mice were individually transferred to an open-topped cage, in a light-, temperature-, and humidity-controlled chamber. Then, mice were tethered to a patch cord (0.57 NA, 400 μm diameter, Doric) with freely rotary joint (Doric) for maximum freedom during movement. After a minimum 3 d of acclimation to tethering, recording was started. The excitation light was provided by a 430 nm LED. To minimize photobleaching during long-term recording experiments, the total laser power delivered into the SCN was adjusted to be as low as possible (10 to 20 μW), and the laser was only turned on for 10 s at intervals of 5 min. Emissions were detected with filters at 460–500 nm (CFP) and 528–556 nm (YFP) using a four-port mini-cube (Doric) and acquired separately using two femtowatt photoreceivers (model 2151, Newport). The FRET ratio (YFP/CFP) was calculated based on the CFP and YFP values recorded during the central 8 s of each 10-s measurement window. Independence (no interference) of FRET recording from ambient light conditions was verified using enucleated mice. We also confirmed that simultaneous and opposing changes in CFP (donor) and YFP (acceptor) intensity were counted as FRET in our photometry recording. At the end of experiments, mice were processed for post-hoc histological evaluation. We eliminated animals where the fiber tip was off target.

### Video-based locomotor activity measurement

Movements of mice were video recorded to assess circadian locomotor activity of mice during fiber photometry. The videos were captured under infrared light using an infrared CCD camera (JN-335CP, JIN). Video images of the entire cage were recorded at a resolution of 360 × 270 pixels. The area of the mouse was roughly 3000 pixels. Images were collected at a frequency of 30 Hz. To reduce computation time, video recordings were down-sampled to 30 frames per min. To automatically find the positions of mice from video images, we first binarized images using the 3000th highest values of frames as thresholds. Then, we clustered binarized pixels using DBSCAN (Scikit-learn library, eps = 5, min_samples = 50), and the largest cluster was defined as the area of animal. The centroid of this animal area was used as a reference point. Locomotor activity was defined as the distance between the reference points in sequential frames. The data were binned into 5-min epochs for tracing circadian locomotor activity.

### SCN slice FRET measurement

A Millicell membrane containing a cultured SCN slice was transferred to a glass bottom (0.1 mm thick) dish, filled with medium. Time lapse FRET imaging was performed essentially as described previously^42,43^. Briefly, we used a two-photon excitation laser scanning incubator microscope (LCV110-MPE, Olympus) equipped with 25×/1.05 NA water-immersion objective lens (XLPLN 25XWMP2, Olympus) and an InSight DeepSee Laser (Spectra Physics). CFP was excited at 840 nm. An IR cut filter (32BA750RIF), three dichroic mirrors (SDM505, SDM570, RDM445XL), and two emission filters (BA460–500 for CFP and BA520–560 for YFP) (Olympus) were used. All recordings were performed every 20 min. Acquired images were analyzed with MATLAB (Mathworks) and the FRET efficiency values of ROI in the image sequence were exported into Excel (Microsoft).

### Sleep recording and analysis

EEG and EMG were recorded by implanted telemetry transmitters (HD-X02, Data Sciences International). For the placement of EEG wires, two small burr holes were made at AP, +1.50 mm, ML, +1.50 mm and AP, −1.50 mm, ML, −1.50 mm from the bregma under general anaesthesia. The ends of the wires were stripped, and one lead was placed through each burr hole so as to touch, but not penetrate, the dura. For EMG, two other wires from the transmitter were placed in the neck/trapezius muscle, each separated from the other by 5 mm. Animals were allowed at least 3-4 weeks to recover from surgery. The EEG and EMG waveform data were acquired using Ponemah software (v.6.41, Data Sciences International) and analyzed using SleepSign software (v.3.3, Kissei Comtec). The data were binned into 10-s epochs and classified into three vigilance stages (wakefulness, NREM, and REM sleep) on the basis of standard criteria for rodent sleep^50–52^. Wakefulness was scored based on the presence of locomotor activity, high spectrum power of EMG, and low spectrum power of delta (1–4 Hz) frequency EEG. NREM sleep was classified based on low EMG spectrum power and high spectrum power of delta frequency EEG. REM sleep was defined as low EMG spectrum power and theta (4–8 Hz) dominant EEG. Semiautomatic sleep scoring was visually inspected and corrected when appropriate.

### AAV preparation and infection for *Gz* reexpression

pAAV-Gz-P2A-mCherry was created by cloning the full-length coding sequence of the mouse *Gnaz* (NM_010311) to the pAAV-human synapsin-1 promoter (hSyn)-P2A-mCherry vector. On the other hand, pAAV-mCherry only encoded mCherry under the promoter hSyn. AAV vectors were packaged in-house based on a published method^48^ in an AAV9 virus serotype. The titers of AAVs were approximately 3.0 × 10^13^ GC/ml, measured by qPCR.

Virus microinjection was performed as previously described^53^ using *Gnaz*^−/−^ mice. Mice were mounted on a stereotaxic micromanipulator (Kopf) under anesthesia. AAV viral vectors were bilaterally injected into the SCN (120 nl/side; coordinates relative to bregma: AP, −0.4 mm, DV, −5.70 mm, ML, ±0.1 mm) at a flow rate of 50 nl min^−1^ using a glass microcapillary pipette, operated through an automated microsyringe pump (KD Scientific Legato 130). The capillary was left in place for 10 min and slowly withdrawn over 5 min. After surgery, mice were returned to the test cages and maintained in LD for recovery and re-entrainment of locomotor activity rhythms. Following proper re-entrainment, mice were released into DD. At the end of the experiments, sites of infection were histologically confirmed, and unsuccessfully targeted animals were excluded from analysis.

For infection to organotypic SCN slices, we applied AAV (2 µl per slice) onto the surface of cultured SCN slices. After 8 h incubation, the culture medium was changed, and time lapse FRET measurements were performed until the end of the experiment.

### Quantitative measures of behavioral activity rhythms

Actogram, activity profile, and χ^2^ periodogram analyses in Fig. 2a and Fig. 4 were performed on ClockLab software (Actimetrics)^54^. The free-running period (τ), total activity level, activity bout length (α), siesta, and rest bout length (ρ) were measured based on animal behaviors in a 14-d interval taken 7 days after the start of the DD condition. The τ was determined by χ^2^ periodogram. The activity onset and offset phases, identified using ClockLab software, were verified by visual inspection (Actimetrics). Activity duration (α) was measured as the time difference between the activity onset and offset of the cycle. Rest duration (ρ) was the time difference between the offset of one cycle and the onset of the next cycle^22,30,31^. Video-based locomotor activity records in Fig. 1 and Fig. 2d were analyzed in the same manner, except for τ, which was analyzed using FFT-NLLS via BioDare2 (biodare2.ed.ac.uk)^55,56^. This is consistent with the τ analysis of FRET ratio rhythms from the same mice. Circadian locomotor/FRET activity patterns of *Gnaz*^−/−^ mice were analyzed based on a 7-d interval taken 7 days after DD.

### Quantitative measures of FRET ratio rhythms

The τ, amplitude, and rhythmicity of FRET ratio rhythms in SCN slice cultures were analyzed using FFT-NLLS (biodare2.ed.ac.uk). Rhythmicity was measured as the reciprocal of relative amplitude error (RAE^-1^). Acrophase was defined as the time interval between the mid-timepoint of the rising phase and the mid-timepoint of the decreasing phase in each cycle. Similarly, the nadir phase length (in Fig. 2f, j) was determined as the interval between the midpoint of the decreasing phase and the midpoint of the subsequent rising phase. On the other hand, the nadir-bottom bout length examined in Fig. 5 was determined as the interval between the point at which the FRET ratio decreased to 20% of its amplitude during the decreasing phase and the point at which it increased to 20% of its amplitude during the subsequent rising phase.

The τ, acrophase, and nadir phase length of FRET ratio rhythms in freely moving mice were determined as described for slice cultures. In all experiments, CT was defined based on the τ of the locomotor activity rhythm, with CT12 being the onset of circadian locomotor activity. The magnitude of a light pulse (30 min, 200 lx)-induced phase shift was quantified as the time difference between regression lines of activity onset before and after the light application^27^. When the data were presented as double-plots in Fig. 1b and Fig. 2f, FRET ratio values were detrended and baseline corrected. For the circular plot in Fig. 1d, events in which mice exhibited circadian FRET and locomotor activity above half of each daily average were extracted, and these events were plotted in Rayleigh format using Oriana 4 software (Kovacs Computer Services)^26^. To determine the 50% ascending or descending time (Time 50%) of FRET and locomotor activity (in Fig. 1e), sigmoidal dose–response curves with variable slope, *Y* = Bottom + (Top – Bottom)/(1 + 10^(log^ ^Time50%-^ *^X^*^)^ ^HillSlope^) were fitted to the data points surrounding CT12 (CT10–CT14) and CT0 (CT22–CT2) using GraphPad Prism software.

### RNA-seq

SCN samples (3 slices per pool) were lysed into ice-cold 20 mM Tris-HCl (pH 7.5) buffer containing 150 mM NaCl, 5 mM MgCl_2_, 1 mM dithiothreitol, and 1% Triton X-100. After digesting genome DNA with DNase I (25 U ml^-1^, Thermo Fisher Scientific), lysates were centrifuged at 20,000 × *g* and the supernatant was processed for library generation as previously described^54^. Total RNA extracts (0.2 μg per sample), purified using Direct-zol RNA Microprep Kit (Zymo Research), were subjected to library construction using SEQuoia Express Stranded RNA Library Prep Kit (Bio-Rad) and sequenced on NovaSeq X plus. Paired-end reads (150 bp) were mapped onto the mouse genome (GRCm38/mm10) using STAR (version 2.7.0), sorted and indexed using SAMtools (version 1.10), and quantified using fp-count (available at https://github.com/ingolia-lab/RiboSeq). For filtering rRNA/tRNA, STAR was used with rRNA and tRNA annotations downloaded from the UCSC table browser. Any RNAs with a read count of 0 in one or more samples were removed, and 23,206 of the total 30,728 annotated RNAs passed this inclusion criterion. To identify transcripts consistently upregulated or downregulated with respect to Gz expression in both the Gz^+/+^ versus Gz^−/−^ comparisons and the Gz rescue versus no rescue comparisons, a two-way group comparison was undertaken using DESeq2 (version 1.44.0). Genes with adjusted P value of < 0.05 were considered as differentially expressed (DEGs). GO enrichment was calculated with GOrilla online tool (http://cbl-gorilla.cs.technion.ac.il/). The complete list of enriched GO terms is available in Supplementary Data 1.

### Luciferase-based cAMP monitoring in organotypic SCN slices

We applied AAV (2 μl; AAV-hSyn-Okiluc-aCT, 3.0 × 10^13^ GC/ml)^24^ onto the surface of cultured organotypic SCN slices from *Gz*^+/+^ and *Gz*^-/-;^ mice. More than 14 days after the viral infection, we started recording. _D_-Luciferin (1 mM) was included in culture medium. Luminescence was measured using a dish-type luminometer (Kronos-Dio, ATTO) maintained at 35°C. Recording was performed for 2 min for each dish at 20-min intervals. The raw data were smoothed using a 1-h moving average and further detrended by subtracting a 24 h running average.

### Optical manipulation of cAMP in the SCN in vivo

*Slc32a1-Cre* mice received injections of AAV-hSyn-flex-bPAC vector^24^ or AAV-hSyn-flex-mCherry into the SCN (3.0 × 10^13^ GC/ml, 120 nl/side, bilateral injection). A flexible fiber-optic cable was connected to an optical fiber implanted in individual mice as described for fiber photometry, and the cable was attached to an LED light source (470 nm; 5 mW at the tip of the fiber; Doric). Stimulation was provided as 20 ms pulses of LED light at a constant frequency of 10 Hz using a pulse stimulator (STOmk-2, BRC Bio Research Center), as previously described^24^. Recording of behavior was performed using video tracking as described above for fiber photometry.

## QUANTIFICATION AND STATISTICAL ANALYSIS

The α, ρ, and τ lengths of behavioral rhythms were determined using ClockLab software (Acti-metrics). We used χ^2^ periodogram for the determination of τ. The values for period, amplitude, and rhythmicity (RAE^-1^) in SCN slices were determined using the FFT-NLLS function of the online BioDare2 suite (https://www.biodare.ed.ac.uk). EEG and EMG waveform data were acquired using Ponemah software (v.6.41, Data Sciences International) and analyzed using SleepSign software (v.3.3, Kissei Comtec). We utilized GraphPad Prism 10 for statistical analysis and data plotting. The statistical details of each experiment, including the statistical tests used, the exact value of *n*, and what *n* represents, are described in each figure legend. For circular statistics, Rayleigh’s uniformity test was performed using Oriana 4 software (Kovacs Computer Services).

## SUPPLEMENTAL INFORMATION

Supplemental information is available for this paper.

## ACKNOWLEDGMENTS

We thank members of the M.D. laboratory, Drs Takahito Miyake and Ms. Kanako Takakura at Kyoto University Live Imaging Center for technical assistance and discussion. We thank Dr William J. Schwartz for critical reading of the manuscript. This work was supported in part by research grants from the Ministry of Education, Culture, Sports, Science and Technology of Japan (22H04987, 24H02306), the Basis for Supporting Innovative Drug Discovery and Life Science Research program of the Japan Agency for Medical Research and Development (JP21am0101092), SRF, the ABiS (22H04926), and the Astellas Foundation for Research on Metabolic Disorders. Y.T. is a recipient of the JSPS Research Fellowship for Young Scientists.

## AUTHOR CONTRIBUTIONS

M.D. conceived the project; M.D. and Y.T. designed the research with M.M., T.H. and H.O.; Y.T. performed experiments in collaboration with Yuto F., T.M., S.K., G.S., A.H., X.S., D.O., Yoshihisa F., and E.H.; M.D., T.M., and Y.T. wrote the paper with input from all authors.

---

**Supplementary Fig. 1.**
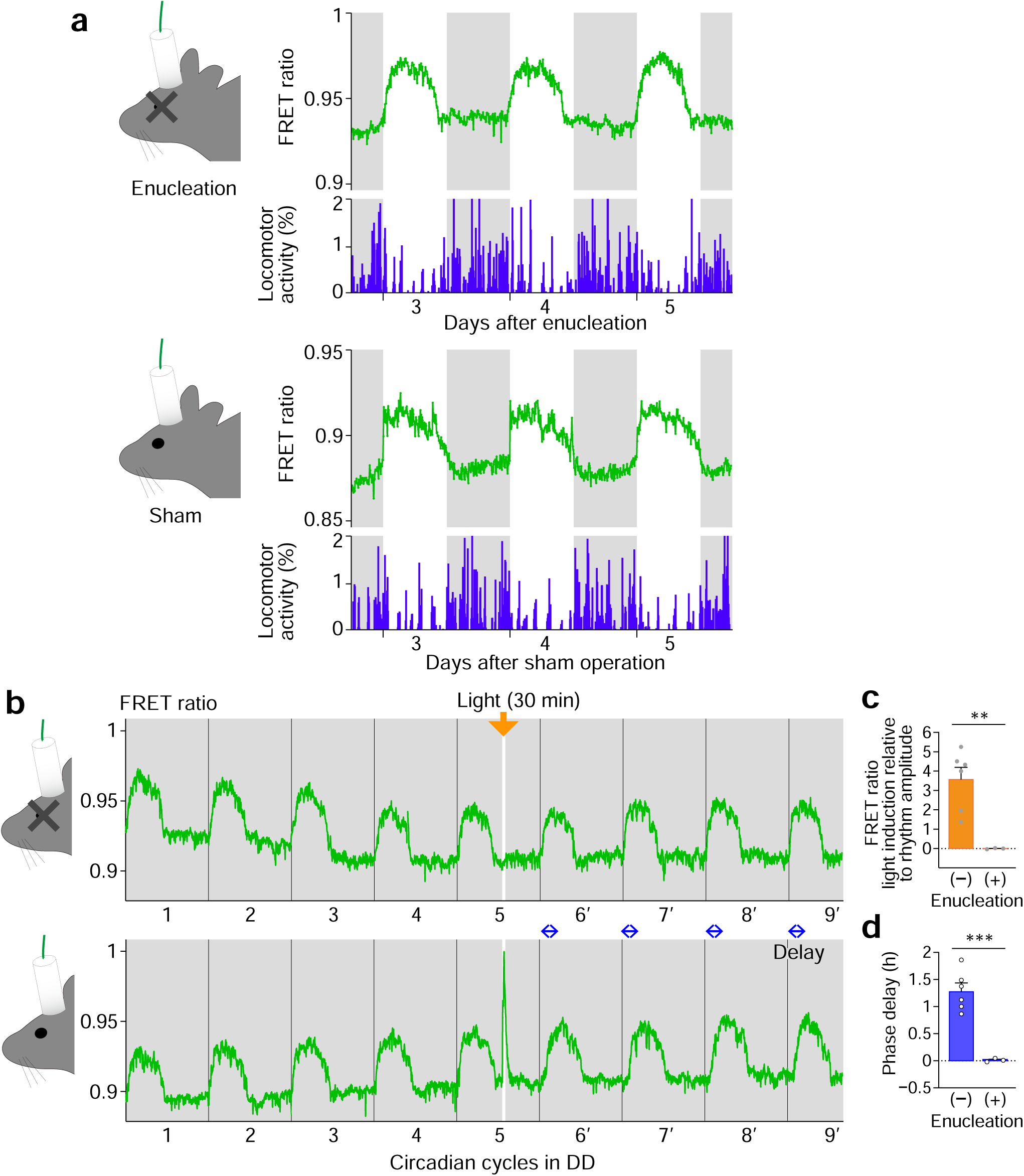
Depletion of ambient light-dependent increase in FRET ratio in enucleated mice. **a** Representative SCN PKA FRET activity profiles of eye enucleated mice and sham operated mice, maintained in LD. Video-monitored locomotor activity profiles are indicated in parallel. Note that acute elevation of SCN FRET ratio in the dark–light transition was absent after enucleation. **b** Representative SCN PKA FRET activity profiles of enucleated mice and sham operated mice maintained in DD followed by a brief light exposure at CT14. **c** Quantification of light induction of FRET ratio, relative to circadian oscillation amplitude. **d** Quantification of phase delay of FRET ratio rhythm. *n* = 3 mice for enucleation; *n* = 6, control. \*\*\**P* < 0.001, \*\**P* < 0.01. Statistics, unpaired Student’s *t*-test.

**Supplementary Fig. 2.**
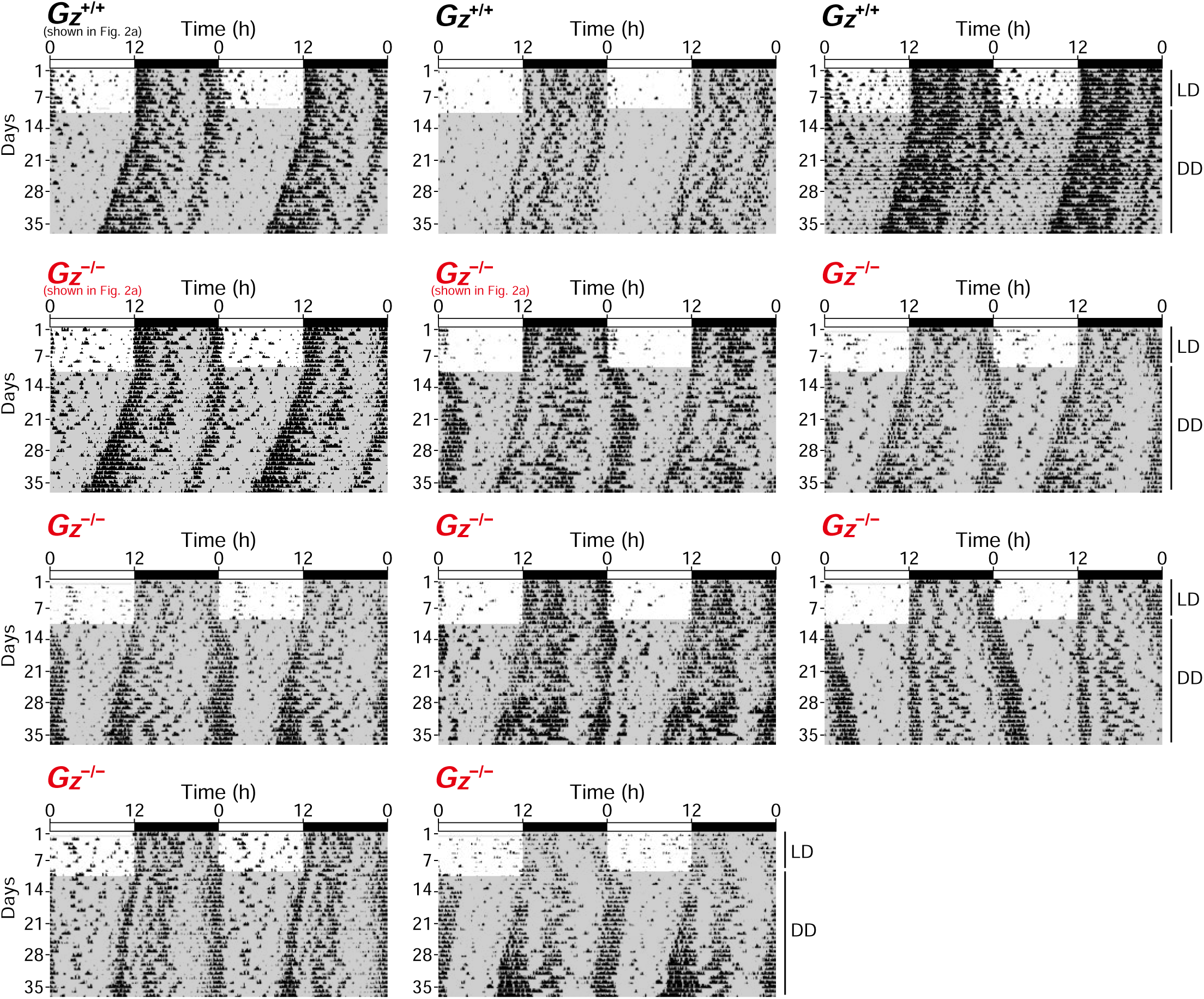
Representative locomotor activity records of C57BL/6J-backcrossed *G*z^+/+^ and *G*z^−/−^mice, related to. Fig. 2a. Mice were housed in LD and then transferred to DD. Periods of darkness are indicated by grey backgrounds. Data are shown in double-plotted format. Each horizontal line represents 48 h; the second 24-h period is plotted to the right and below the first.

**Supplementary Fig. 3.**
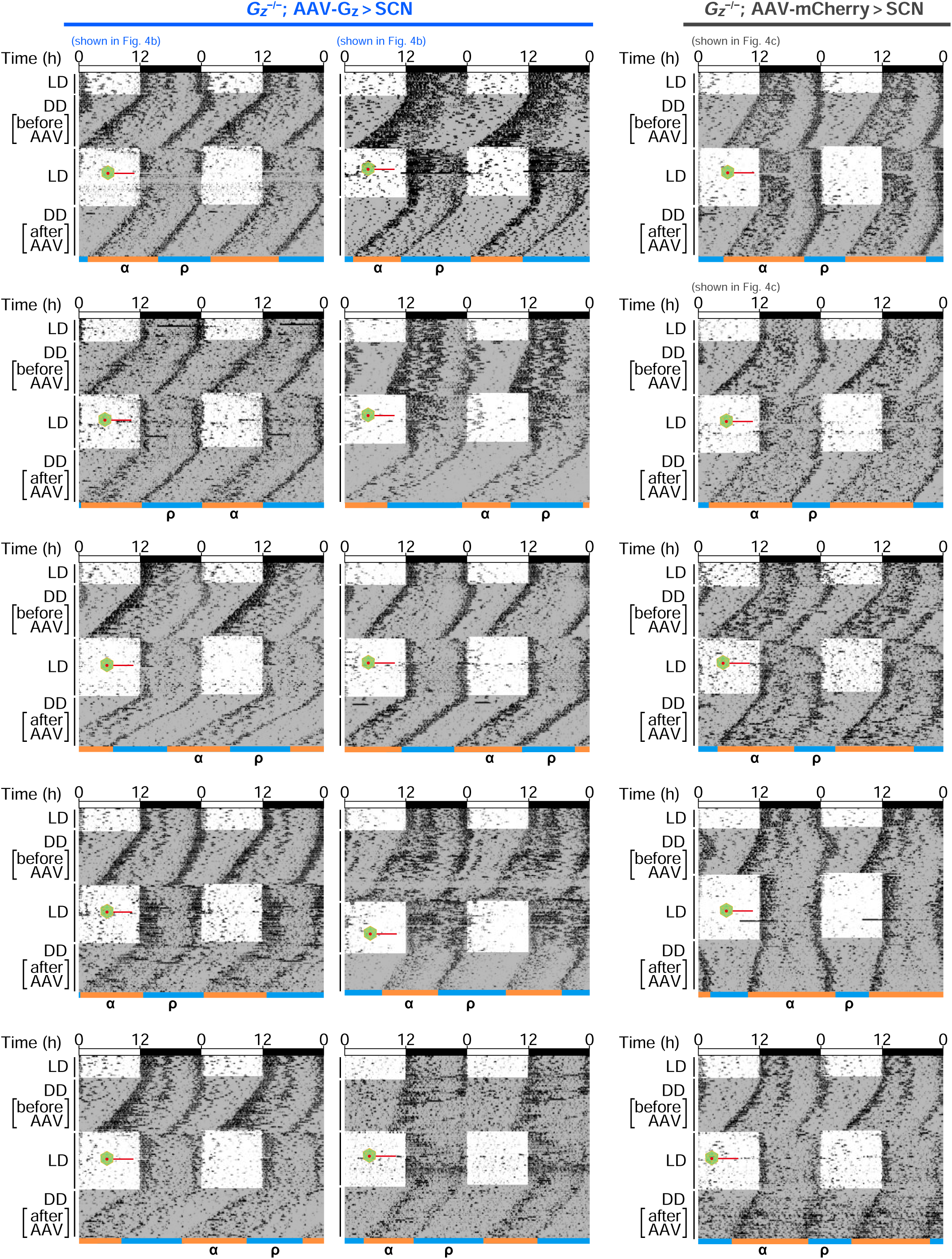
Representative locomotor activity records of AAV-*G*z rescue experiments in. Fig. 4. Mice were housed in LD and then transferred to DD to monitor basal behavioral activity profiles before AAV-*Gz* rescue. Injection of AAV-Gz-2A-mCherry or its control AAV-mCherry to the SCN of *Gz*^-/-;^ mice was then performed after reentrainment to LD cycles. Red lines indicate the time of operation for viral injection. After surgery, mice were returned to the test cages and maintained in LD for additional 17–19 d for recovery and to allow for viral expression. Mice were then released into DD to monitor their spontaneous locomotor activity profiles. Note that viral introduction of *Gz*, but not *mCherry*, rescues normal rest length (ρ) in *Gz*^-/-;^ mice.

**Supplementary Fig. 4.**
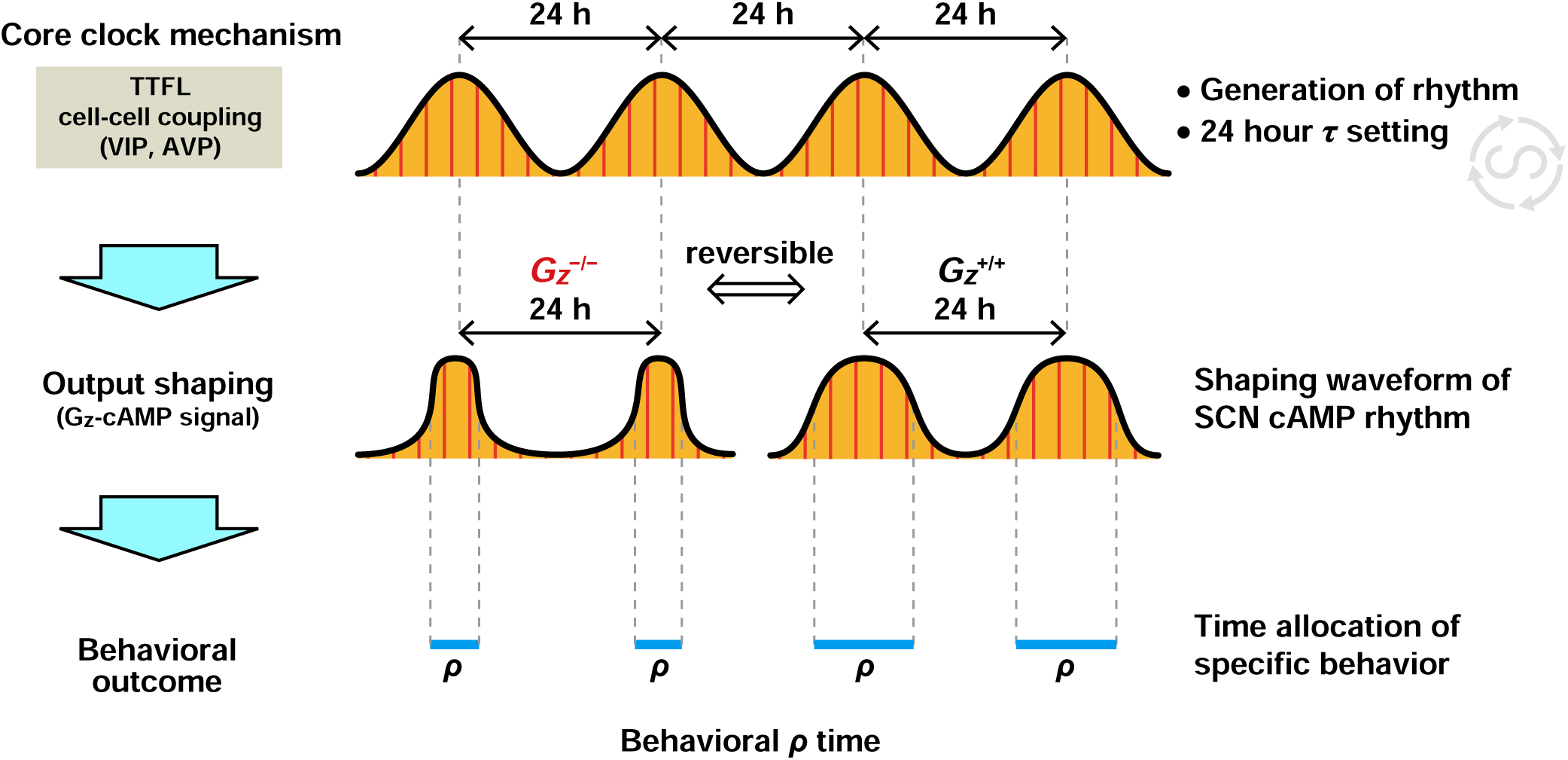
A model depicting a role for Gz in circadian clock system. Gz is not necessary for either rhythm generation or circadian period determination. Rather Gz has a specific contribution to output shaping. Specifically, Gz contributes to shaping the acrophase of circadian cAMP-PKA activity rhythm in the SCN and thereby defines the time and duration of circadian rest timing (*ρ*) of the organism. The shape of output rhythm govering *ρ* can be reversibly modified by Gz.

